# Humans parsimoniously represent auditory sequences by pruning and completing the underlying network structure

**DOI:** 10.1101/2022.05.19.492659

**Authors:** Lucas Benjamin, Ana Fló, Fosca Al Roumi, Ghislaine Dehaene-Lambertz

## Abstract

Successive auditory inputs are rarely independent, their relationships ranging from local transitions between elements to hierarchical and nested representations. In many situations, humans retrieve these dependencies even from limited datasets. However, this learning at multiple scale levels is poorly understood. Here We used the formalism proposed by network science to study the representation of local and higher order structures, and their interaction, in auditory sequences. We show that human adults exhibited biases in their perception of local transitions between elements, which made them sensitive to high-order structures such as network communities. This behavior is consistent with the creation of a parsimonious simplified model from the evidence they receive, achieved by pruning and completing relationships between network elements. This observation suggests that the brain does not rely on exact memories but compressed representations of the world. Moreover, this bias can be analytically modeled by a memory/efficiency trade-off. This model correctly account for previous findings, including local transition probabilities as well as high order network structures, unifying statistical learning across scales. We finally propose putative brain implementations of such bias.

## Introduction

**T**o interact efficiently with their environment, humans have to learn how to structure its complexity. In fact, far from being random, the sensory inputs we face are highly interdependent and often follow an underlying hidden structure that the brain tries to capture from the incomplete or noisy input it receives. For instance, ***Tenenbaum et al. (2011***), proposed that learning implies building the simpler underlying relational model which can explain the data. Indeed, evidence suggests that humans can infer structures from data at different scales, ranging from local statistics on consecutive items (***Saffran et al., 1996***) to local and global statistical dependencies across sequences of notes (***Bekinschtein et al., 2009***; ***Basirat et al., 2014***) or more high order and abstract relationships such as pattern repetition (***Barascud et al., 2016***), hierarchical patterns and nested structures (***Dehaene et al., 2015***), networks (***Schapiro et al., 2013***; ***Garvert et al., 2017***) and rules (***Maheu et al., 2020***).

At first, the extraction of local regularities in auditory streams was proposed as a major mechanism to structure the input, available from an early age since ***Saffran et al. (1996***) showed that 8-month-old infants could use transition probabilities - P(A|B) - between syllables to extract words from a monotonous stream with no other available cues. Since then, the sensitivity of humans to local dependencies has been robustly demonstrated in the auditory and visual domain (***Fiser and Aslin, 2002***), without the focus of attention (***Batterink and Paller, 2019***; ***Benjamin et al., 2021***; ***Batterink and Choi, 2021***) and even in asleep neonates (***Fló et al., 2022***; ***Benjamin et al., 2022***). Moreover, it is not limited to adjacent elements but can be extended to non-adjacent syllables - P(A|XC) - that could account for non-adjacent dependencies in language (***Peña et al., 2002***).

However, the computation of transition probabilities (TP) between adjacent - P(A|B) - and non-adjacent elements - P(A|XC) - seems too limited to allow the extraction of higher-order properties without an infinite memory that the human brain does not have. Network science - an emerging interdisciplinary field - thus proposed a different description to characterize more complex streams ***Lynn and Bassett (2020***).In this framework, a stream of stimuli corresponds to a random walk in the associated probabilistic network. Several studies used this network approach to investigate how humans encode visual sequential information (***Garvert et al., 2017***; ***Mark et al., 2020***). Shapiro and colleagues (***Schapiro et al., 2013***) tested human adults with a network consisting of three communities (i.e. sets of nodes densely connected with each other and poorly connected with the rest of the graph - (***Newman, 2003***)) where transitions between all elements were equiprobable (each node had the same degree). This community structure is an extreme version of the communities and clustering properties often found in real-life networks, whether social, biological or phonological (***Girvan and Newman, 2002***; ***Karuza et al., 2016***). The authors observed that subjects discriminated transitions between communities from those within communities. Since local properties (TP) were not informative, this result revealed participants’ sensitivity to higher-order properties not covered by local probabilistic models. Recently, Lynn and colleagues replicated a similar effect with a probabilistic sequential response task (***Lynn et al., 2020***). They presented subjects with sequences of visual stimuli that followed a random walk into a network composed of three communities. After each stimulus, subjects were asked to press one or two computer keys, and their reaction time was measured as a proxy of the predictability of the stimulus. To explain the response pattern, the authors proposed an analytical model that optimizes the trade-off between accuracy and computational complexity by minimizing the free energy function. This model allows taking into account the probability of memory errors in the computation of the transition probabilities (TP) between the elements of the stream. From now on, we will refer to this model as the Free-Energy Minimization Model (FEMM : Model E, explained below).

In this paper, we aim to merge these two lines of research and validate a model that can explain how humans learn when local and high-order relations are simultaneously present in sequences generated from noisy or incomplete structures. Moreover, we propose that adults do not encode the exact input but a condensed version based on the generalization of the underlying structure. To this end, we took advantage of the community network framework and adapted it to expose adult participants to sequences of sounds that followed a random walk in a network as in the studies described above (***Schapiro et al., 2013***; ***Lynn et al., 2020***), but using sparse communities, that is with missing transitions between elements of the same community (see Fig 1). This design allows investigating whether participants are able to complete the network according to the high-order structure or if, on the contrary, they rely on local transitions and reject impossible transitions independently of the high-order structure. In other words, after training with an incomplete network, if new (“unheard”) transitions are presented, are participants more willing to accept them if they belong to the community (i.e., within-community transitions) than if they occur between communities? Moreover, while several papers have studied network learning in the visual domain (***Schapiro et al., 2013***; ***Karuza et al., 2019***; ***Lynn et al., 2020***), to our knowledge, it has never been tested in the auditory domain. Given the sophisticated auditory sequence processing capacities observed in humans compared to other primates (***Dehaene et al., 2015***) and their potential importance in language acquisition, it is important to investigate the auditory domain.

**Figure 1.**
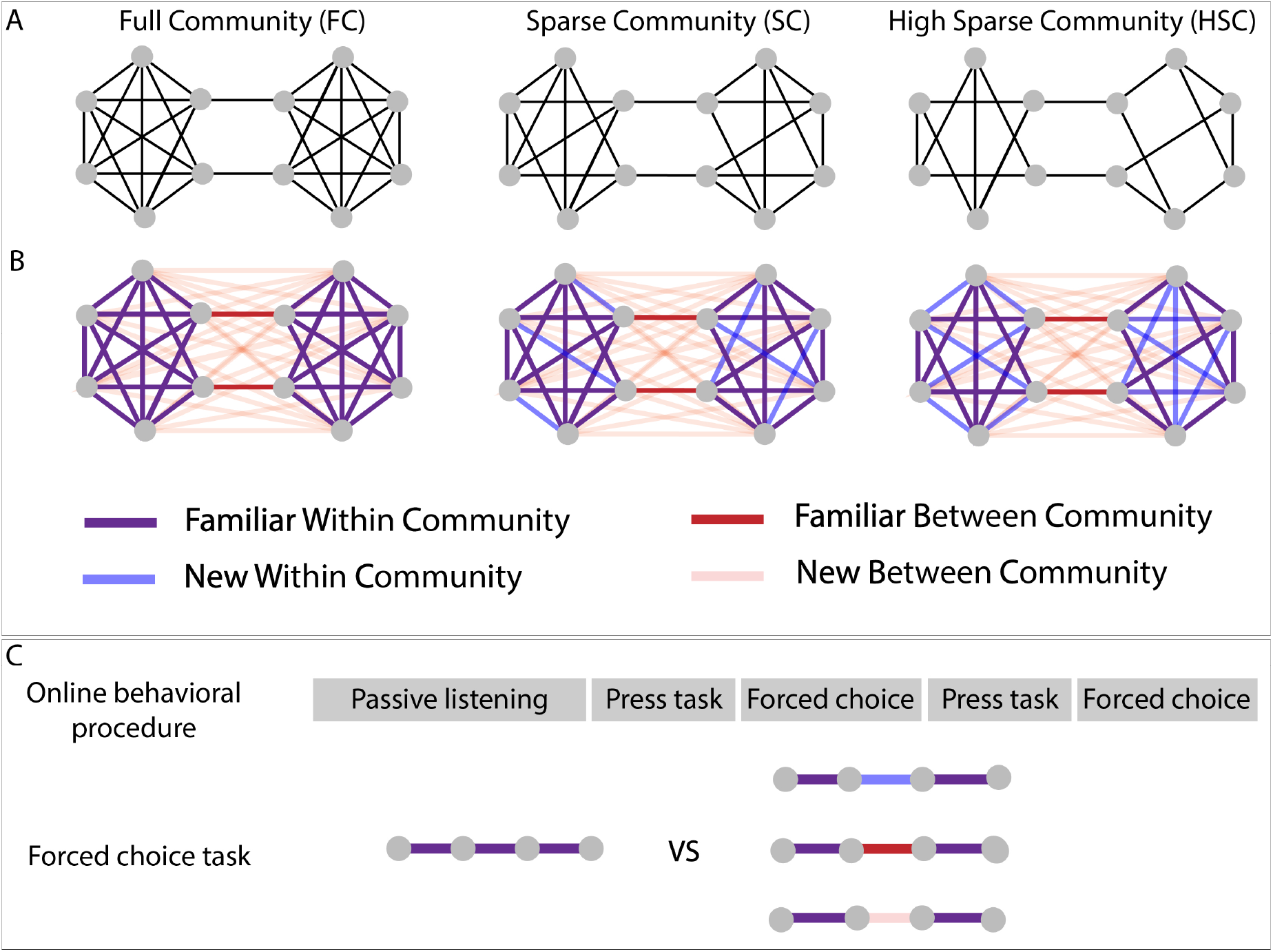
A: Graph structure to which adult subjects were exposed in three different experiments B: Graph design with color coded conditions. Blue and pink lines represent transitions that have never been presented during the stream presentation but only during the forced-choice task. C: Test procedure used for behavioral testing. In the press task phase, participants had to press a key when they felt there was a natural break in the sequence. In the forced-choice task, they had to chose between the two quadruplets, the most congruent with the sequence they had heard. In the proposed pair, one was always a Familiar Within Condition transition (purple transitions), and the other, one of the three other conditions

We tested three different experimental designs, using both pure tones and syllables. After a training phase of four minutes, during which participants were asked to attentively listen to a sequence of sounds, we asked them to press a key when they felt there was a natural break in the sequence during two additional minutes of active listening. Finally, we assessed what was retained in memory by asking them to choose between two isolated quadruplets, the most congruent quadruplet with what they had heard before (Two-Forced Choice Task). This phase allowed us to present previously unheard transitions (“new transitions”) and to study whether participants were able to generalize the network structure (Fig 1). The first tested experimental network - Full Community - was composed of two communities of 6 elements each, with all nodes within a community connected with each other (except two nodes at the border of the community to keep an equal degree for each node). During the forced-choice task, we tested transitions within the community already presented during the stream (*Familiar Within Community transitions*) against transitions between communities (*Familiar Between Community transitions*) and unheard transitions crossing communities (*New Between Community transitions*). We expected here to replicate in the auditory domain Lynn et al’s results obtained in the visual domain and to assessed memory retention using two forced choice recall. The second design - Sparse Community - was similar to the Full Community except that one possible edge for each node was removed, allowing to test a new condition during the forced-choice task: *New Within Community transitions* corresponding to the unheard transitions within communities. Finally, in a third design, we tested even sparser communities - High Sparse Community - obtained by removing two possible edges per node from the Full Community. The performances in these two “sparse” designs, relatively to the full-community design, are crucial to investigate the participants’ underlying representations of the sequences.

Many studies on sequence learning proposed different and not always compatible ad-hoc models to account for their results. Therefore we compared the participants’ behavior to the predictions of those different models proposed in the stream processing and graph learning literature (fig2). Specifically, we designed paradigms that maximize the differences between the predictions of the different models.

**Figure 2.**
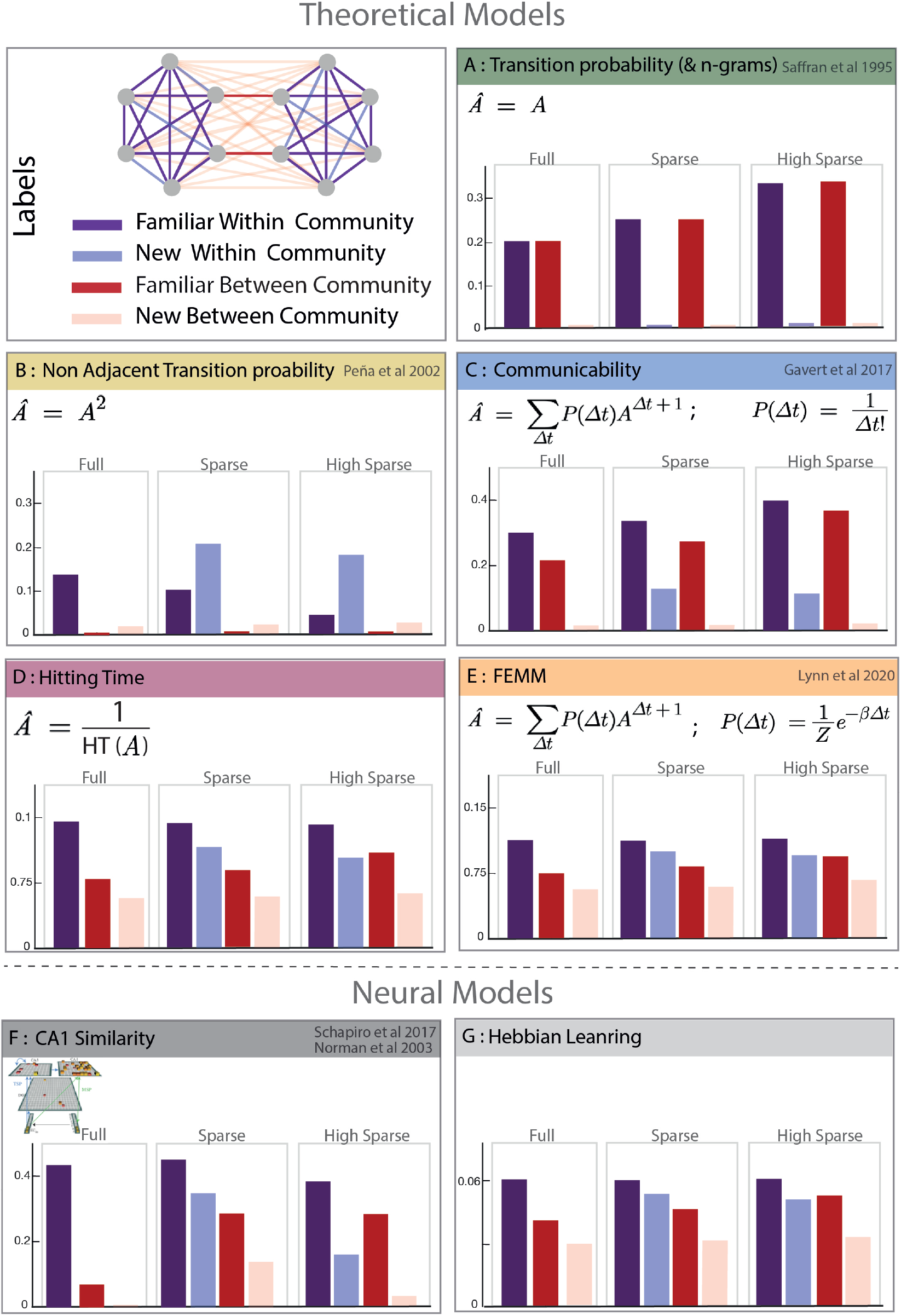
Model description and predictions for the three paradigms tested. For each model, we computed the value predicted in each condition in the Full, Sparse, and High Sparse paradigms. Although the models are partially correlated, they differ in their prediction about the congruency of New Within Community transitions (light blue). Models A, B, C, D, and E are theoretical metrics over the graph structure that predict more or less familiarity with the different types of transitions. Model F and G are biologically plausible neural encoding of those metrics. The box colors correspond to the conditions labeled in the top-left panel.

- *Model A: Transition Probabilities (TP) and Ngrams* : Local transitions between consecutive elements - P(A|B) - have been proposed as an efficient learning mechanism to structure streams of input. We tested the limits of this simple local learning computation in the presence of a high-order structure. Ngrams are similar to TP but take into account n previous items in the computation of the transition. For example for trigrams, *P* (*E*_*t*_|*E*_*t*−1_*E*_*t*−2_*E*_*t*−2_). Note that because our designs are random walks into Markovian networks, the transition probabilities and Ngram models are identical. *P* (*E*_*t*_ |*E*_*t*−1_*E*_*t*−2_*E*_*t*−2_) = *P* (*E*_*t*_|*E*_*t*−1_). In our designs, there are two kinds of local transitions: Familiar transitions and New transitions (TP = 0). Since the TP calculation does not consider community structure, new transitions should be rejected by participants regardless of their relation with respect to communities (New Within Communities = New Between Communities).
- *Model B: Non adjacent TP*: This metric is similar to the transition probabilities but on non consecutive items *P* (*E*_*t*_| *XE*_*t*−2_). We included it in our analysis because several studies have shown human sensitivity to such properties in streams (***Peña et al., 2002***).
- *Model C: Graph Communicability*: This model comes from the network science literature and computes the relative proximity between nodes in the network, making it sensitive to cluster-like structures like communities. Interestingly, a recent study shows that this measure correlates with fMRI data (***Garvert et al., 2017***), suggesting a potential relevance in human cognition.
- *Model D: Hitting Time*: This metric is also coming from network science. It computes the average number of edges needed (path length) to move from one node to another during a random walk. Similar to Communicability (model C) and FEMM (model E), it measures a ‘proximity’ between nodes in a network. To make it more comparable with the other models we computed its inverse value.
- *Model E: Free Energy Minimization Model (FEMM)*: This model, recently proposed by ***Lynn et al. (2020***) to account for community sensitivity by humans, is a trade-off between accuracy and computational complexity. It can be explained by memory errors while computing Transition Probability (TP) between elements in a stream. Participants exposed to a stream of elements reinforce the association between element *i* and *i-1*. However, errors in this process may lead participants to sometimes bind element *i* with element *i-2, i-3, i-4* with a decreasing probability (for a full description of the model see ***Lynn et al. (2020***)). Mathematically, the distribution of the error size that minimizes the free energy function is a decreasing exponential (Boltzmann distribution). Therefore, the estimated mental model of transition probability is biased compared to the streams’ objective transition probabilities enabling participants to encode high-order structure. In more detail, the mental model is a linear combination of the transition probability matrix (A) and non adjacent transition probabilities of every order (*A*^Δ*t*^) with a weight of *P* (Δ*t*) where Δ*t* is the order of non adjacency (or size of the memory error, ie Δ*t* = *n* corresponds to *P* (*E*_*t*_|*X*… *XE*_*t*−*n*_)). The estimated model can then be written as

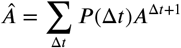

with

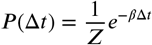

where A is the transition probability matrix of the graph. *β* was previously estimated to 0.06 in a comparable task with human adults (***Lynn et al., 2020***). We therefore first used this value to test this model on our behavioral data and later confirmed this estimation with our data (see SI).

Although the different models are partially correlated with each other, they give different predictions about network pruning and completion,i.e. about the perceived familiarity of New Within Community and Familiar Between Community transitions (Fig. 2). FEMM and Hitting Time models predict that participants should better detect New Between Community than New Within Community transitions, which is also partly the case for the Communicability model, but not for the TP model. The TP model predicts that participants should only distinguish “heard” (familiar) transitions from “unheard” (new) transitions. The high similarity between FEMM, Hitting Time, and Communicability models is not surprising as they all describe the same property of the network: proximity between nodes. Intuitively, items from the same community will appear closer together than items from different communities, even if the two nodes are not connected. In fact, FEMM and Communicability are mathematically very close but with a different decay (exponential vs. factorial).

In addition to those theoretical models, we considered two putative brain implementations using biologically realistic neural networks :

- *Model F: Hippocampus CA1 similarity*: This neural network aims to reproduce the hippocampus structure (***Norman and O’Reilly, 2003***), which is often described as a key structure in statistical and structure learning (***Schapiro et al., 2017, 2016***; ***Henin et al., 2021***). We compute here the similarity in CA1 layer as it has been proposed to capture community-like structures in previous studies (***Schapiro et al., 2017***).
- *Model G: Hebbian learning with decay*: Hebbian learning is a biologically plausible cortical implementation of associative learning. Some neurons fire specifically to some objects in the environment. When two of those neurons co-fire, the pair is reinforced. The implementation of transition probabilities learning has been proposed as a cortical mechanism based on this principle. Here we adapted this idea to implement FEMM instead of TP, specifically by adding a temporal exponential decay in the probability of a neuron firing after a stimulus’s presentation. When the exponential decay has the same *β* parameter as the FEMM, the results of the FEMM and the Hebbian learning with decay are highly correlated.

## Results

### Human Behavior

All participants were exposed to a stream of either tones or syllables adhering to one of three possible graphs (Fig. 1A and 1B). After a 4-mn-familiarization period, they were instructed to press the spacebar when they had the impression of stimuli change (active listening task). Then they were presented with a two-forced-choice task to decide between two sequences which one was more congruent with the structure of the stream they listened to (Fig. 1C). No differences were found between the experiments using tones and syllables. Thus, data were merged in the following analyses.

#### Key presses distribution during active listening

Fig 3 top row shows the normalized distribution probability of key presses after a transition, using a kernel approach (see methods for detailed computation). In all three paradigms (each corresponding to a graph in Fig 1), there was a significant increase in key presses after transitions Between Community compared to transitions Within Community (all p<0.05 are indicated by bold lines), revealing that participants were sensitive to switching between sound communities. Full Community and Sparse community designs showed a similar effect size, while the High Sparse Community design elicited a significantly smaller effect size. Unpaired t-tests every ms in [-0.1, 2.750] s window, contrasting the Full Community vs. High Sparse Community, show a significant difference between 1.024 s and 2.631 s post-transition (p<0.05 Bonferroni corrected). Similarly, Sparse Community vs. High Sparse Community differed between 0.798 s and 2.548 s (p<0.05 Bonferroni corrected).

**Figure 3.**
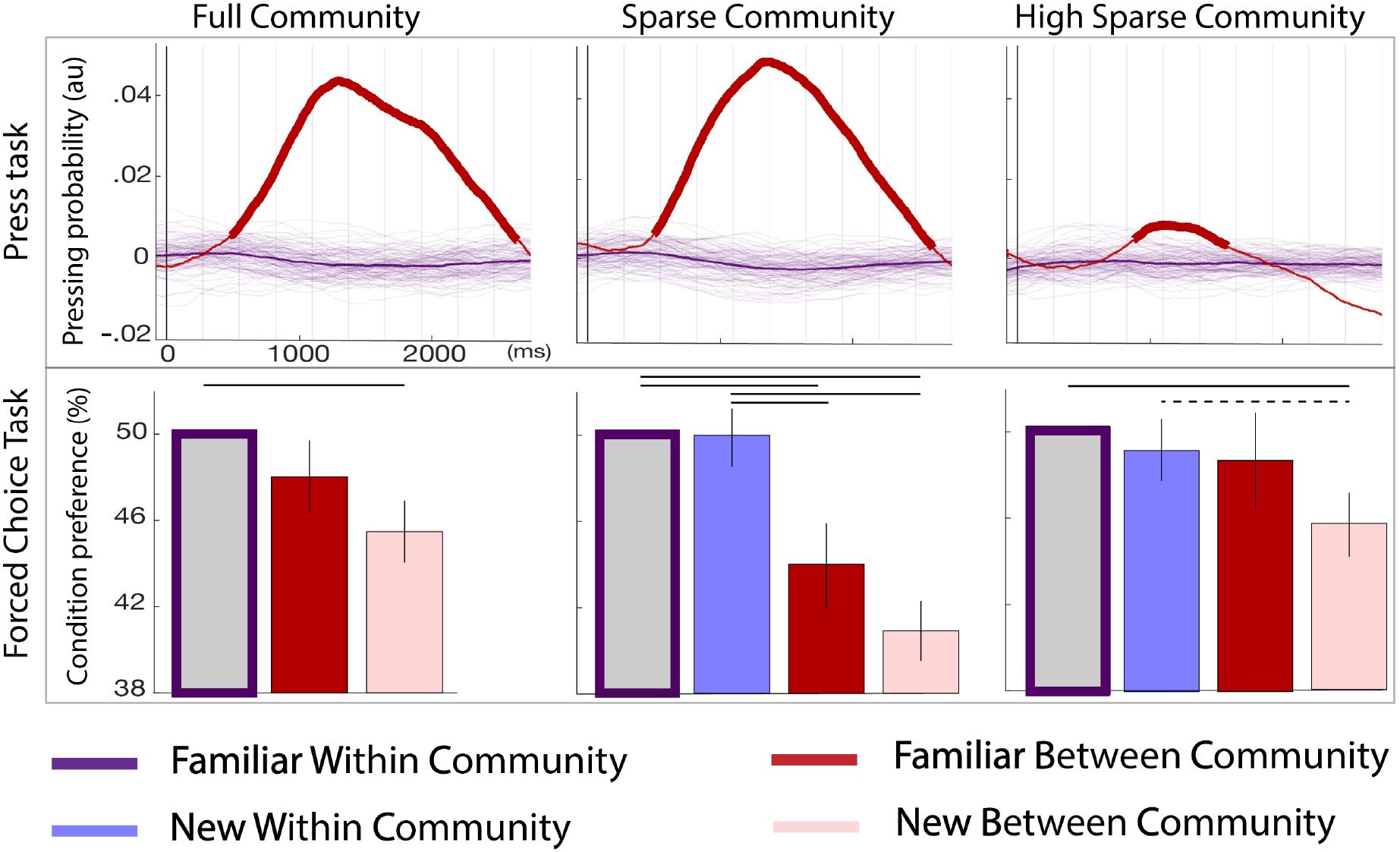
Behavioral results for the three paradigms. Top panel: parsing probability during the active listening phase (distribution of key presses after the offset of a given transition) purple lines: Familiar Within Community transitions, red line: Familiar Between Community transitions. Thin purple lines each represent a bootstrap occurrence of the parsing probability for the Familiar Within Community transition. The bold red line indicates the time-points where there was a significant increase of parsing probability after a Familiar Between Community transition compared to a Familiar Within Community transition. Bottom panel: Percentage of responses for each condition during the forced-choice task. By convention, the Familiar Within Community condition, which was presented at each trial, is set to 50%. The plain lines above the boxes indicate significant differences between conditions (pval<0.05 FDRcor) and the dotted lines marginal significance (pval < 0.05 uncorr)

#### Two-forced-choice task

To estimate subjects’ memory encoding of the transitions and test impossible transitions in the graph, we presented a two-forced-choice task in which subjects had to choose between a Familiar Within Community transition and one of the other conditions. Fig 2 shows the percentage of preference for each condition. 50% represents chance level while <50% score represents a preference for the Familiar Within Community transition compared to the other conditions. If participants encoded familiar transitions, they should reject any new transitions more than the Familiar Between Community transition. On the other hand, if they encoded the underlying structure of the communities, we expected them not to recognize the novelty of the New Within Community transitions and reject the two inter-community conditions (familiar and new). First, participants significantly rejected the New Between Community transitions in each experiment (ps<0.01 FDR), although the Familiar Between Community transitions were only significantly rejected for the Sparse Community design (p<0.01 FDR). In contrast, the New Within Community transitions were never rejected (i.e., were confounded with familiar transitions) in the Sparse Community and the High Sparse Community. We also estimated the interaction between each condition. In the Sparse Community paradigm, the New Within Community transitions were significantly less rejected than the Familiar Between Community transitions (p<0.05 FDR) and the New Between Community transitions (p<0.01 FDR). In the High Sparse paradigm, the difference between the New Within Community and the New Between Community transition was only marginally significant (Uncorr p = 0.046). In other words, participants were sensitive to the underlying network structure and naturally completed the graph.

### Adequacy of the models to explain participants’ behavior

#### Correlation between human data and theoretical model predictions

To estimate the adequacy of the models to explain the behavioral data, we pooled together the three studies and used bootstrapping with replacement to estimate the correlation between the data and the model predictions. Fig 4A shows the correlations’ distribution between the data and each model (presented on the diagonal) and between pairs of models. We estimated the significance of the correlation strength between the data and model *i* or *j* by counting the percentage of occurrences in which model *i* had a stronger correlation with the data than model *j*. All models were significantly correlated with the data (all p< 0.01 FDRcorr), with a correlation strength following the order FEMM ≈ Hitting Time > Communicability > Non-adjacent Tps ≈ TPs (Fig 4B). Note that the FEM and Hitting Time models had the best correlation with the data (81%) and were significantly better than the other models (p<0.05 FDR).

**Figure 4.**
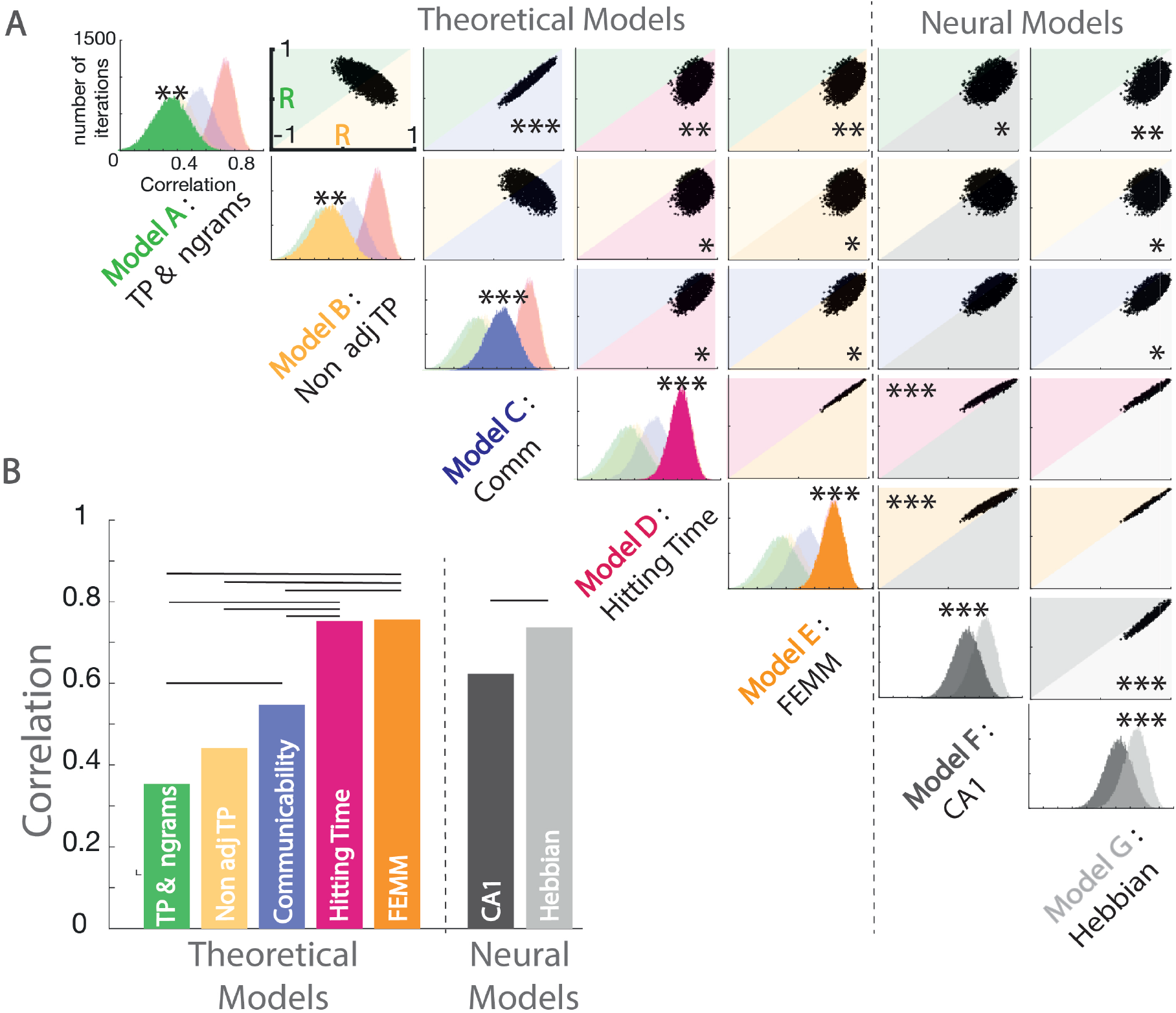
A: Estimation of the correlation of the human data with each model by bootstrap re-sampling. The diagonal of the matrix displays the distribution of correlations between each model and the data computed separately for the theoretical models (A to E) and the neural models (F G). Each panel of the diagonal presents the same data (A to E, then F G), but the data are colored to highlight a specific model to facilitate the comparison between models. For each pair, the significance between models (indicated by stars) is estimated by counting the number of bootstrap occurrences for which one model was more correlated with the data than the other. We plotted this bootstrap as a cloud of dots in the Correlation with Model1 x Correlation with Model2 subspace. Significance is then represented by the percentage of dots above the diagonal. Models with similar predictions display a line style cloud of dots aligned along the diagonal. B: Summary of the correlations between each model and the behavioral data. Plain lines above the boxes indicate the significant differences between models

#### Correlation between human data and neural model predictions

As the FEMM computation and the Hitting Time were the best theoretical models, we translated them into a realistic biological architecture using Hebbian rules. We estimated this implementation on a 50000 items long stream for each paradigm. The correlation between the analytical computation and the Hebbian learning implementation exceeds 99%. Using the same bootstrap approach, we compared this Hebbian approach with a neural network reproducing hippocampus architecture proposed by ***Norman and O’Reilly (2003***). Both models were significantly correlated with the data and with the other. However, the Hebbian implementation of FEMM was significantly better than the hippocampus model (Fig 4).

## Discussion

### Transition probabilities between elements of the sequence is biased by the structure of the underlying generative network

Our results show that human adults do not encode transition probabilities objectively when familiarized with a stream of sounds. Instead, they seem to have a systematic bias to complete the transitions within a community suggesting a subjective internal representation that differs from the objective distribution of the transitions they heard. This behavior is compatible with two proposed theoretical models: the Free Energy Minimization Model (FEMM) and the Hitting Time.

The high agreement between the Free Energy Minimization Model and the data we observed, suggests that the bias can be analytically estimated using the FEMM *Â* = Σ_Δt_*P*(Δ*t*)*A*^Δ*t*+1^ with 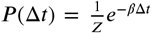. ***Lynn et al. (2020***) proposed that this bias can be explained by memory errors while trying to recall the previous item of the stream during the TP computation.

The bias in the encoding of Transition Probability between successive elements enabled the extraction and encoding of high-order structures in graphs, i.e., a community structure. We can distinguish two distinct bias effects: First, the *pruning* of familiar transitions that do not obey the community structure (i.e., Familiar Between Community transitions are rejected). Second, the *completion* of the structure by over-generalizing new transitions when those are compatible with the high-order structure (i.e. New Within Community transitions are accepted). These perceptive biases lead to a more parsimonious internal representation of graphs.

### Putative brain implementation of such computation

We showed that computations of transition probabilities were biased in humans and analytically characterized this bias as an optimal trade-off between accuracy and computational complexity. Indeed, perfect accuracy in the encoding would result in being insensitive to the high order structure while, on the other hand, too low accuracy would result in no learning at all. We also tested putative brain implementations that could account for such a computation in the human brain. Specifically, we tested to which extent two previously described mechanisms could account for our results: Cortical Hebbian learning and Hippocampus episodic memory.

Hebbian learning is a very simple mechanism consisting of the reinforcement of pairs co-occurring in a signal. It has been proposed as a learning mechanism for statistical learning tasks (***Endress and Johnson, 2021***). Here, we minimally modified it as described above to introduce the bias in TP computation. Such learning could be directly implemented in cortical regions by synaptic plasticity induced by learning.

Another hypothesis focuses on the hippocampus. In fact, several authors propose that statistical learning and graph learning might be represented as the construction of an abstract map of relational knowledge, analogous to topographic maps (***Garvert et al., 2017***; ***Constantinescu et al., 2016***) that are known to involve the hippocampus. A recent experimental study (***Henin et al., 2021***), showed that when exposed to statistically organized auditory or visual streams, the hippocampus activity measured with ECoG exhibited a cluster-like behavior, all elements belonging to the same group being similarly encoded. Using the community paradigm with fMRI, ***Schapiro et al. (2016***) also reported an increased pattern similarity in the hippocampus for elements belonging to the same community. Another piece of evidence comes from modeling the hippocampus activity in different statistical learning tasks (***Schapiro et al., 2017***). In this study, the authors used a neural model mimicking the hippocampus architecture and trained it on different statistical learning tasks including, community structure learning. They showed that the pattern of activity in CA1 might account for both pair learning (episodic memory) and community structure learning. This result might be partially consistent with two mechanisms observed in the hippocampus: pattern completion (i.e. the similarity of the neural representations of close stimuli increases, which allows generalization) and pattern separation (i.e. the similarity of neural representation of close stimuli decreases, to disambiguate them) (***Yassa and Stark, 2011***; ***Liu et al., 2016***; ***Bakker et al., 2008***). In our experiment, despite similar acoustical distance between stimuli, tones from the same community might benefit from a pattern completion phenomenon due to an increase in the similarity of their neural representation when they belong to the same community. On the other hand, the pruning effect described here might also correspond to a pattern separation mechanism decreasing the similarity of the neural representation of tones belonging to different communities. The completion and pruning effect behaviorally reported in our experiment might thus rely on these two mechanisms operated by the hippocampus.

We compared here those two hypotheses (Hebbian and hippocampal learning) by comparing the predictions of the hippocampus model used by ***Schapiro et al. (2017***) and a biased Hebbian learning approach. We showed that both models fit the data very well, with a slightly better result for the Hebbian learning approach. Since we only have behavioral results, it is difficult to conclude on the brain mechanisms involved, especially since recent work has proposed the joint use of cortical and hippocampal learning in similar tasks (***Whittington et al., 2020***). In any case, the agreement between the behavioral data and the two brain models shows that the FEMM model (an analytical model), does not only explain behavioral data but also has biologically valid candidates.

### A general model of sequence learning

Statistical learning has been proposed as a powerful general learning mechanism that might be particularly useful in language acquisition in order to extract words from the speech stream (***Saffran et al., 1996***). However, the exact model explaining statistical learning remains under-specified: What is computed remains unclear (***Henin et al., 2021***; ***Fló et al., 2022***) and authors often tailored the computation to suit the paradigm (Transition probabilities in some studies, non-adjacent or backward transition probabilities in others, biased transitions probabilities in network studies…). We argue that the Free Energy Minimization Model (FEMM) is a more general model that, beyond explaining community separation, as shown above, can also account for results traditionally explained by the computation of local transitional probabilities and those that require the computation of long distance dependencies. Indeed, the first-order approximation of the Free Energy Minimization Model corresponds to the objective Transition Probabilities model (*Â*_0_ see SI). Thus, the predictions of the FEMM model are the same as those of the transition probability model in many tasks, notably in classical speech segmentation experiments, where a drop in TP signals word edges (***Saffran et al., 1996***).

Another part of the statistical learning literature focuses on AxC structures, in which the first syllable of a triplet predicts the last syllable (***Peña et al., 2002***; ***Buiatti et al., 2009***; ***Endress and Johnson, 2021***; ***Kabdebon et al., 2015***; ***Marchetto and Bonatti, 2015***). The computation of first-order TPs is insufficient to solve this task, which requires the encoding of non-adjacent TPs. However, a bias estimation of TPs following the FEMM is sensitive to non-adjacent dependencies and can explain the emergence of AxC structures. Additionally, as previous papers and our results show, the FEMM can also explain subjects’ behavior in different kinds of network learning (***Schapiro et al., 2013***; ***Karuza et al., 2016***; ***Lynn et al., 2020***). Lynn and colleagues (***Lynn et al., 2020***) interpret the FEMM as errors in the associations between elements, whose probability decays with the distance between associated elements. We proposed that implementing the TPs computation through Hebbian learning with a firing decay results in a comparable computation to the Free Energy model.

Finally, a similar Hebbian learning approach enables to explain the sensitivity to backward TP reported in the literature (***Pelucchi et al., 2009***; ***Endress and Johnson, 2021***). A similar idea has recently been proposed by ***Endress and Johnson (2021***).

However, the authors did not refer to free energy optimum and did not provide an analytical approach. Instead, they proposed a Hebbian learning rule with the same idea of mixing TP with non-adjacent TP (which corresponds to a second order approximation of the Free Energy Minimization Model that we propose here, see *Â*_1_ in SI). Like we do here, they argued that this mechanism could account for results currently explained by different models in the literature. Thus, the FEMM model and its putative neural implementation through Hebbian rules unifies different proposals concerning statistical learning on the one hand and network learning results on the other hand, under a common principle.

### Information compression and stream complexity

Our results showed that humans have a biased subjective representation of first-order transition probabilities compared to the actual transition probabilities, enabling them to learn high-order structure in the underlying graph and overgeneralize transitions that they never experienced. What is the advantage of such a computational bias for human cognition? We distinguish three main advantages.

1. *Higher-order structures and generalization can be relevant information to learn*. Unlike random networks, many real-world networks have transitivity properties (***Girvan and Newman, 2002***; ***Newman, 2003, 2006***) - if A is connected to B and B to C, there is a high chance for A and C to be connected (a friend of my friend is likely to be my friend). Thus, when building an abstract network representation of elements dependencies, it is relevant to be sensitive to this type of dependencies.
2. *Overgeneralizing enables fast learning*. Overgeneralizing means accepting transitions congruent with the structure even before they appear in the stream. Thus for short exposures, the estimation of the Free Energy Minimization Model is closer to the real transition probability matrix than the estimation of the transition probability based on the input. Because many real-world networks are highly transitive, this over-generalization is a good prediction of the underlying network. For example, in the case of the Full Community paradigm, the FEMM estimation is a better estimator of the actual Transition probability matrix than the estimation of transition probability for streams shorter than 150 elements approximately. If the amount of information is limited, the FEMM bias is optimal and faster than the unbiased learning of TPs. This fast learning might be of importance, for example, for language acquisition, given that human infants are exposed to a limited amount of speech, the classical poverty of the stimulus argument.
3. *The extraction of high-order structures enables information compression*. Because of the computational cost in a biological system and the pressure on memory to encode long sequences, compressing information is a major advantage. In a community paradigm such as the ones tested here, the stream can be reduced to the extreme to a binary sequence with a certain probability of changing between communities A and B (Fig 5). Instead of remembering all the transitions of the stream, it is sufficient to remember the community label and probability of transition between communities. For the sparse community paradigm, the probability of staying inside a community is 92%, then the average number of consecutive elements of the same community is 12.5. Therefore, the average K-complexity of a stream is divided by 6.25 thanks to compression (see Fig 5). Recent data (***Planton et al., 2021***; ***Al Roumi et al., 2020***; ***Sablé-Meyer et al., 2021, 2022***) showed that humans’ performances were highly sensitive to stimuli compressibility, arguing for a compressed encoding of inputs. Thus, the extraction of high-order structures could be at the basis of a human abstract compressed language of thought (***Fodor, ????***) that will enable a more efficient manipulation of complex information. In this same line, a recent study using a graph perspective (***Whittington et al., 2020***) proposes that the representation of the abstract relational structure in a stream of elements (the underlying graph) and the mapping between node and stimuli identity could be factorized. This factorized representation would enable an abstract representation that allows fast learning in scenarios sharing the same graph structure by translating the structure to new nodes-stimuli mappings. Thus, our present results suggesting that humans form a compressed representation of the relational graph structure, add to this line of ideas arguing for a simplified world model created by the human brain.

**Figure 5.**
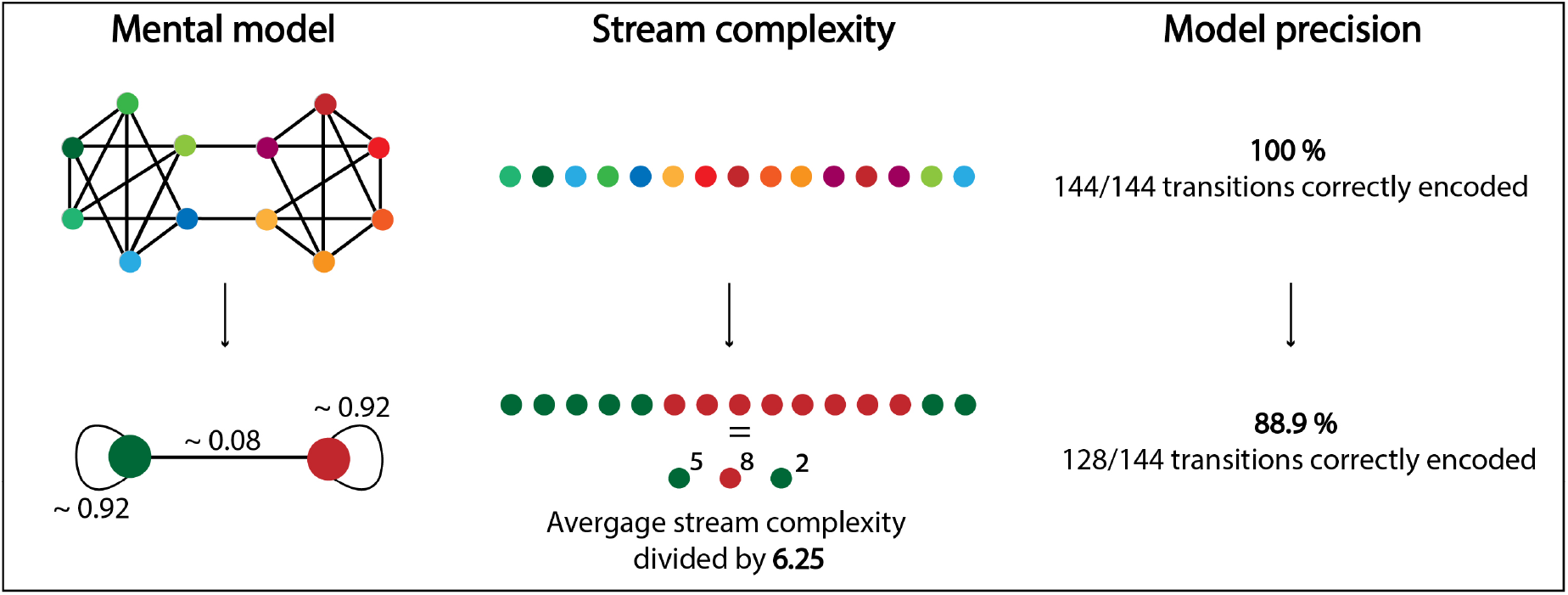
Compression of the Sparse community graph is efficient as it divides the average K-Complexity of a stream by 6.25 while keeping 89% of the information about transitions correctly encoded.

Finally, the human sensitivity to community is in line with Simon’s postulate that the complexity of a system can only be handled thanks to its hierarchical nearly decomposable property (***Simon, 1962***). In other words, a complex structure is no more than the sparse assembly of less complex dense substructures. Here we propose empirical arguments by demonstrating that humans are sensitive to the decomposition of a complex network into two simpler sub-networks.

### Methodological remarks

In this study, we used two different metrics. The press bar task during attentive listening showed high sensitivity, but it only allowed to test within vs between community transitions during learning and thus assess for pruning effect and clustering. On the other hand, the forced-choice task on the isolated quadruplets allowed testing for all conditions after learning. However, this second metric has low sensitivity, and only a few trials can be collected, resulting in very high variability in the data. To overcome this, we collected a very large sample of participants online (N=583). A more sensitive method would be needed to explore the brain bases of this kind of paradigms.

### Conclusion

The results shown in this study reveal 1) Community representation in the auditory domain; 2) The persistence of a biased, subjective transition probabilities representation after learning; and most importantly, 3) *pruning* and *completion* effects allowing to build a parsimonious representation of the underlying network structure. Transition probabilities are thus not exactly encoded by the participants but biased in a way that can be predicted by the Free Energy Minimization computation. The same model might explain human sensitivity to local and high-level regularities without the need for specific models for each task. More research is needed to characterize how and where such computations take place in the human brain and how this bias varies across individuals and with development. However, Hebbian rules in the cortex and eventually the hippocampus might be plausible candidates for a biological implementation of this analytical model. Finally, finding appropriate metrics to cluster graphs is a current research topic in applied mathematics (***Newman, 2006***). Thus we believe that understanding the cognitive processes at stake when humans are exposed to such structured networks might provide insight to cognitively and biologically plausible computations.

## Materials and Methods

### Behavioral Task

#### Participants

A total of 727 adults were recruited via social media (424 of which were retributed 2.5$on Prolific platform) with no reported auditory issues or language related troubles. They were assigned to one version of the experiments and instructed to carefully listen for 4 minutes to a nonsense language composed of nonsense words that they had to learn because they would have to answer questions on the words afterward. Participants were either exposed to the Full Community (N= 250), the Sparse Community (N= 249), or the High Sparse Community (N= 228) paradigms with either pure tones or syllables as stimuli.

#### Stimuli

We generated twelve tones of 250ms duration, linearly distributed from 300 to 1800 Hz. Each experiment was composed of 4 minutes of an artificial monotonous stream of concatenated tones without any pause, resulting from a random walk into the tested graph. To avoid any putative acoustical bias, we collected 8 groups of subjects for each paradigm. For each of the 8 groups, we generated a new graph (for FC, only one graph was possible, for SC and HSC we randomly created 8 graphs), a new correspondence between the alphabet of tones and the nodes of the graph and finally new random walks into the graph.

We also collected data sets on the three paradigms with an analog procedure and instructions but using syllables. All speech stimuli were generated with the MBROLA text-to-speech software (***Dutoit et al., 1996***), using French diphones, with a duration of 250 ms, flat intonation and no coarticulation between syllables.

#### Procedure

Participants started with a 4-mn familiarisation phase of exposure to the stream. Then learning was tested with two tasks. First, participants were told that the order of the tones/syllables was not random and that they had to press the spacebar when there was a noticeable change in the tones (or syllables) group used in the stream. Then, they were again exposed to a random walk stream for 2 mn (active listening). Second, they were presented with a two-forced-choice task in which they had to choose between two quadri-elements sequences. They had to choose the most likely sequence, part of the language they learned. The two-forced-choice trials always comprised a Familiar Within Community transition and one representing the other conditions. These conditions were New Within Community transitions, New Between Community transitions, and Familiar Between Community transitions (Fig 1). Participants were exposed to 8 trials of each type (with different sounds each time) except for the New Within Community type, where they were only exposed to 4 trials because, by design, there are only 4 of those transitions in the graphs. Each transition used in the set was presented in both directions (AB and BA). 4 catch trials were also included to control participants’ engagement in the task. These catch trials were two consecutive identical quadruplets that subjects had to detect. The testing part (active listening and forced-choice task) was repeated twice.

#### Data Processing: Active listening

Participants who pressed less than 10, or more than 200, times during the experiments were excluded from further analysis (FC: 52/250; SC: 24/249; HSC: 23/228). A null array of the size of the stream was built and filled with ones at times when participants pressed the space-bar (Dirac impulses). To convert it into a continuous signal, we convoluted it with an exponential window. Then, we epoched this continuous signal from -2.75 to 2.75 seconds after each transition’s offset. Finally, we averaged all the epochs corresponding to the four Familiar Between Community transitions and four out of all Familiar Within Community transitions, and compared them. We repeated this with 1000 random groups of four Familiar Within Community transitions in each subject. By normalizing and averaging across subjects, we were able to estimate the increase of the pressing probability after a Familiar Between Community transition compared to a Familiar Within Community transition at each time point. This method is similar to the kernel approach for estimating probability density from discrete observations.

#### Data Processing: Forced-choice task

Participants that failed on more than 2 catch trials (two identical quadruplets) out of 4 were excluded from further analysis (FC: 35/250; SC: 45/249; HSC: 34/228).For each subject, we computed a percentage of preference for the new transition relative to the Familiar Within Community transition in each condition (i.e., the ratio between the number of trials where the subject chose the new sequence and the total number of trials of this condition). The measure ranges from 0 (pure selection of the Familiar Within Community transition) to 100 (pure selection of the other transition) with a chance level of 50%. We estimated the difference with chance level (50%) and between conditions using paired t-tests. In fig3, we presented the participants’ preference choice with the baseline at 50% in grey to make the comparison between behavioral data and model estimations straightforward. We report the data from the second forced-choice-task session, corresponding to maximum exposure to the streams. For the tone stream, results were similar in the first and second sessions. For the syllable stream, results from the first session were more variable, probably because the task was more difficult when syllables were used rather than tones. The correlation analysis was replicated with each sub-group of data (first vs. second session and tones vs. syllables) (see supplementary material).

### Modelling

#### Theoretical models

For the four models that could be analytically computed from the transition probability matrix (A, B, C, and E), we computed the predictions made by the models for each of our graphs (8 with syllables, 8 with tones). Given *A* the transition matrix of the graph, models were computed using the analytical description:

- *Model A: Transition Probabilities (TP) and Ngrams* : By construction of the Transition matrix, the transition probabilities between nodes are the elements of A.

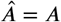
- *Model B: Non adjacent TP*: Non-adjacent TP are computed by taking the square of the transition matrix.

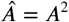
- *Model C: Graph Communicability*:

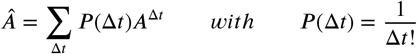 Thus, corresponds to the exponential serie:

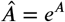

We use Matlab function ‘expm’ to compute this value.
- *Model E: Free Energy Minimization Model (FEMM)*:

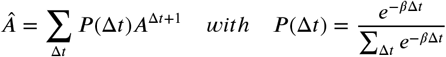

which can be re-written:

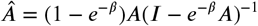

We then computed the average estimate for each of the conditions for each design. Only the Free Energy Minimization Model (Model E) had one free parameter in its equation. To remove this free parameter and make the model more comparable to the others, we used a previously estimated value of ß=0.06 reported in the literature (***Lynn et al., 2020***). To confirm this estimation, we then ran a correlation between our data and the Free Energy Minimization Model for different 10^−15^ and 10^15^ (see SI). The communicability model as described in (***Garvert et al., 2017***) Garvert et al. uses the adjacency matrix. Here we used the transition probability matrix. We believe it is more appropriate to consider the relative weights of each transition and not only its existence or not because a random walk into a weighted graph follows the transition matrix and not the adjacency one. It makes it also more comparable with the other models.
- *Model D: Hitting Time* : For this Model, we approximated its value by creating 50000 items long streams corresponding to each graph and computing the average number of elements between each pair of stimuli. We took the inverse of this value to make it more directly comparable with the other metrics,.

#### Neural Models

- *Model F: CA1 similarity*: We used the neural network and the procedure explained in ***Schapiro et al. (2017***) originally published by ***Norman and O’Reilly (2003***). We did not change any parameter from this original study because our goal was to see how predictable this model was for our paradigms. We trained it 25 times on each of our graph structures (for each paradigm, 25 batches for 8 groups with Syllables and 8 groups with tones: 25*8*2 = 400 replications). We then presented after each training each node as input in isolation and recorded the pattern of activity in the CA1 layer. To estimate the similarity in nodes’ encoding, we computed the correlation between the pattern of activity in CA1 for pairs of elements. We then made predictions on our task by comparing the similarity between two nodes linked by our four types of transitions.
- *Model G: Hebbian Learning with decay*: The aim of this model was to propose an implementation of the FEMM computation with an adaptation of the Hebbian approach proposed for cortical associative learning. To achieve that, we declared a layer of neurons with at least one neuron per node of the graph (it can contain more for generalization to bigger networks). The neurons started firing with an exponential decay corresponding to the FEMM decay for each sound in the sequence. Thus, if another sound was presented before the previous neuron stopped firing, several neurons encoding for different nodes co-fired simultaneously. It biased the estimation of TP between two elements. This co-firing behavior can be computed using Hebbian learning rule to update the weights between the neurons. This weight Matrix is then an estimation of the Free Energy Minimization Model that will converge as the length of the input stream increases. To estimate this model we followed the same procedure as for the Hitting Time. We created 50000 items long streams corresponding to each graph and used those streams as inputs of the neural network. We updated the weigh matrix at each step using Hebbian rule as described before. The weight matrix after the 50000 items was used as an estimation of the model.

#### Model Comparison

To compare models and data, we considered all experimental designs together. To make it comparable with the two-forced-choice data, we normalized each design prediction by the model’s value for Familiar Within Community transitions. We then pooled all data from all designs and estimated the correlation between the data and models’ predictions using 5000 bootstrap re-sampling occurrences. The p-values were estimated by counting the percentage of bootstrap occurrences correlating more with one model compared to another. All the bootstrap occurrences and their correlation with each pair of models are presented in Fig 4B. Each dot represents one bootstrap occurrence. The distribution of these dots below and above the diagonal indicates the comparison between the two models. The scatterplot’s shape shows the correlation, independence, or anticorrelation between two models. This main analysis of data and model comparison has been performed for each subgroup of data (first/second session; tones/syllables). The results in each subgroup are presented as supplementary materials.

## References

This research has received funding from the European Research Council (ERC) under the European Union’s Horizon 2020 research and innovation program (grant agreement No. 695710). We thank Stanislas Dehaene for discussions and remarks during the design of the experiment

## Supplementary Information

### Variation of the beta parameter

As explained in the main text, we used the previously estimated value for *β* = 0.06 (***Lynn et al., 2020***) to remove the free parameter of the model and make it more comparable with the other models of the literature. In order to confirm that this estimation corresponded to our data, we computed the correlation between the subjects’ data and the Free Energy Minimization Model predictions for *β* ranging from 10^−15^ to 10^15^. We smoothed this correlation vector to avoid local variations and found a plateau of high correlation for *β* = [10^−4^; 10^−1^] with a maximum for *β* = 0.049 (Correlation 81%). Similarly, we computed the correlation between the FEMM and the hitting time estimation as a function of *β*. Here again, following the same procedure, we found a plateau of high correlation from *β* = [10^−4^; 10^−1^] with a maximum for *β* = 0.053 (Correlation = 99.3%). The two models can then be considered quasi equivalent with the *β* parameter considered in this paper (0.06).

### Free energy approximation of order n

As described in the main text, FEMM is an infinite weighted sum of Transition Probabilities (*P* (*E*_*t*_|*E*_*t*−1_)), Non Adjacent transition probabilities (*P* (*E*_*t*_|*XE*_*t*−2_)), Non Adjacent transition probabilities of second order (*P* (*E*_*t*_|*XXE*_*t*−3_)) etc… which can be formally written

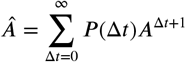

with

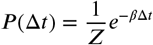

and A the transition probability matrix between all nodes of the graph. An alternative explanation of our data could be that subject compute only TP (*A*) and non adjacent TP (*A*^2^) and combine both evidence later on for decision making. We investigated this, and more generally the nth order approximation of our model, by decomposing *Â* in two parts: the first n elements of the sum and the others. We can then write:

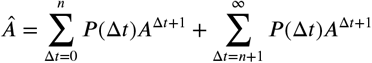

Let’s call *Â*_*n*_ the first n elements of the sum:

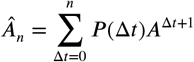

*Â*_*n*_ is then converging toward *Â* as n increases. Note that *A*_0_ is equivalent the computation of transition probability. We estimated the correlation between the behavioral data and *Â*_*n*_ as a function of n ranging from 0 to 15 to estimate which approximation of this infinite sum is enough to correctly represent the data. The correlation between our data and the Free Energy Minimization Model *FEMM* = *Â*_∞_ is 81%. We found that the correlation logarithmically converged toward *Â*_∞_ with 95% of the final value reached for n=4 and 99 % for n=8 (fig S1). The second order approximation including only TP (*A*) and non adjacent TP (*A*^2^) seems then not sufficient to fully explain our data.

### Correlation between human data and model by subgroup

In fig 4, we presented the results of the correlation between human data and the different models when mixing tone and syllables streams together. Moreover, we only showed data from the second two-forced choice task session, corresponding to maximum training. Below we show the same analysis with tone and syllables groups separated for the first and second two-forced choice sessions (Fig S2). The first session in the syllable stream did not show any sensitive pattern. This may be related to participants’ a priori that syllables are ordered in words and that a random walk among syllables contradicts the usual organization of speech, a problem that does not exist with tones. However, the pattern has been replicated three times in the other groups, showing high confidence in this result.

**Figure 6.**
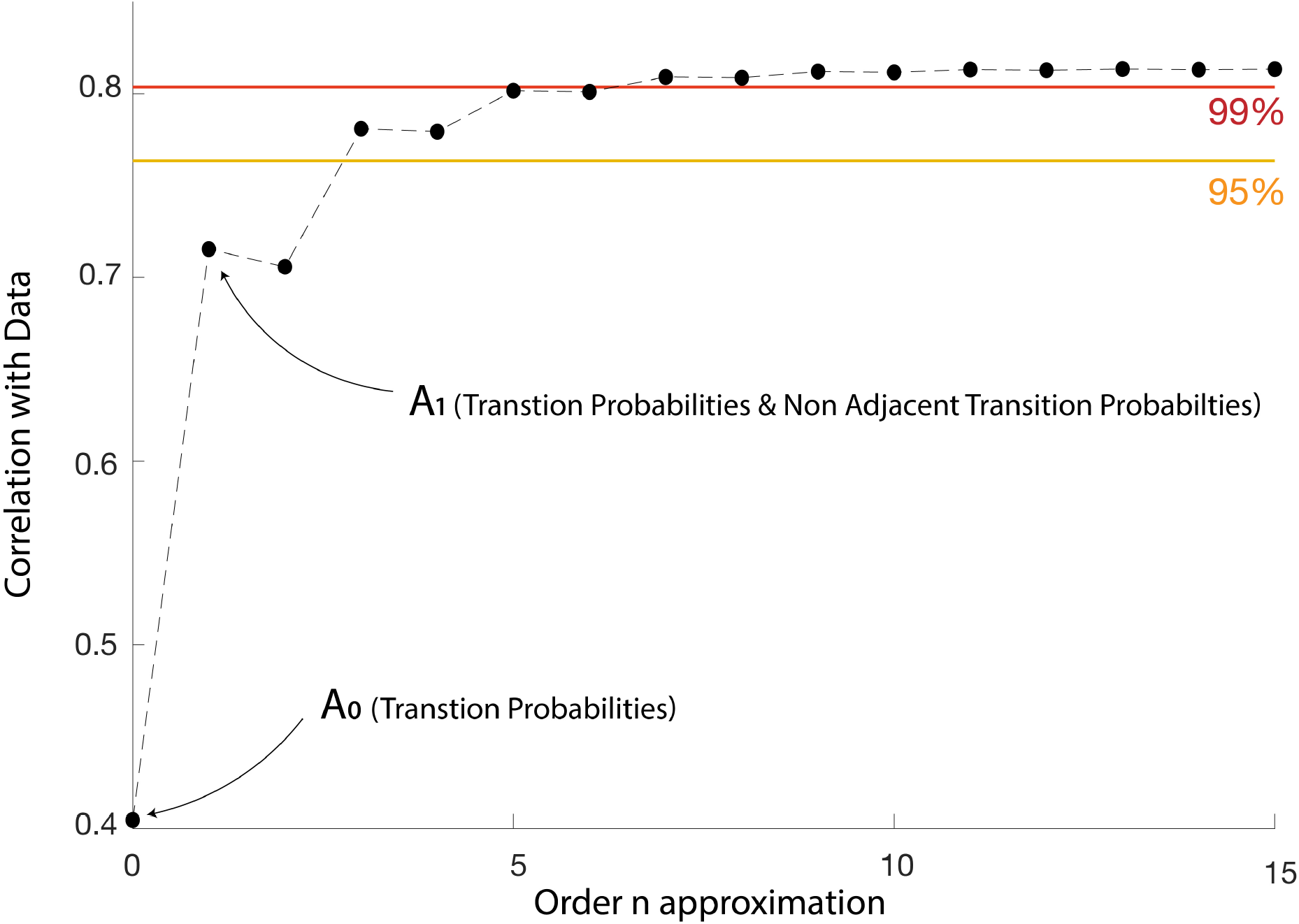
Correlation of *Â*_*n*_ with behavioral data as a function of n. 95% of the maximum correlation is obtained for n=4, 99 % for n=8.

**Figure 7.**
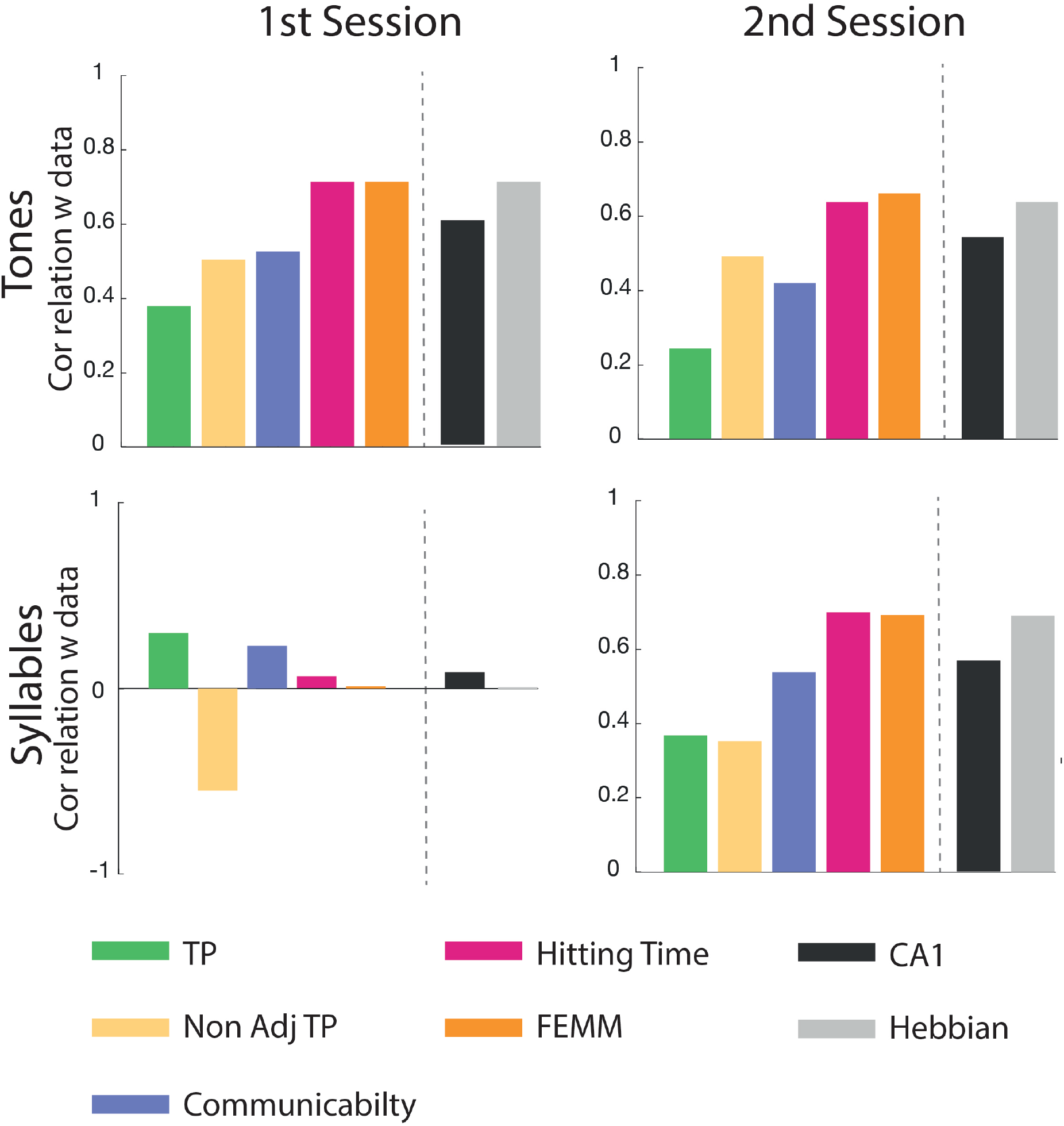
Correlation between models and data divided by test sessions and subgroups (tones and syllables)

